# The anatomical boundary of the rat claustrum

**DOI:** 10.1101/596304

**Authors:** Christopher M Dillingham, Mathias L Mathiasen, Bethany E Frost, Marie AC Lambert, Emma J Bubb, Maciej M Jankowski, John P Aggleton, Shane M O’Mara

## Abstract

The claustrum is a subcortical nucleus that exhibits dense connectivity across the neocortex. Considerable recent progress has been made in establishing its genetic and anatomical characteristics, however a core, contentious issue that regularly presents in the literature pertains to the rostral extent of its anatomical boundary. The present study addressed this issue in the rat brain.

Using a combination of immunohistochemistry and neuronal tract tracing, we have examined the expression profiles of several genes that have previously been identified as exhibiting a differential expression profile in the claustrum relative to the surrounding cortex. The expression profiles of parvalbumin, crystallin mu (Crym), and guanine nucleotide binding protein (G protein), gamma 2 (Gng2) were assessed immunohistochemically alongside, or in combination with cortical anterograde, or retrograde tracer injections. Retrograde neuronal tracer injections into various thalamic nuclei were used to further establish the rostral border of the claustrum.

Expression of all three markers delineated a nuclear boundary that extended considerably (~500 μm) beyond the anterior horn of the neostriatum. Cortical retrograde and anterograde neuronal tracer injections, respectively, revealed distributions of cortically-projecting claustral neurons and cortical efferent inputs to the claustrum that overlapped with the gene marker-derived claustrum boundary. Finally, retrograde tracer injections centred in nucleus reuniens, whilst including the rhomboid, mediodorsal and centromedial nuclei, revealed that insular cortico-thalamic projections encapsulated a claustral area, with strongly diminished cell label, that corresponded to the claustrum.

## Introduction

The claustrum is a highly conserved nucleus that is not only present in all placental species (Baizer et al., 2014) but is also found in *Aves* (Puelles et al. 2016). The claustrum also exhibits genetic characteristics (Mathur et al., 2009; Smith and Alloway, 2010a; Pirone et al., 2012; Hinova-Palova et al., 2014a; b; Sanchez-Vives et al., 2015; Kim et al., 2016; Qadir et al., 2018), and cortical connectivity (Kitanishi and Matsuo, 2016; Patzke et al., 2014; Smith et al., 2014; Smith and Alloway, 2010a; Wang et al., 2017; White et al., 2017; Zingg et al., 2018) that appear to be largely conserved across species (see Buchanan and Johnson, 2011). Progress in understanding the complexities of the rodent claustrum have, however, been hindered by both its irregular shape as well as its small cross-sectional area, factors that have precluded, for instance, effective electrophysiological characterisation. Progress has also been held back by a lack of clarity concerning the extent of its anatomical boundaries, an issue that is seated in the fact that rodents are lisencephalic and, as such, lack a well-defined extreme capsule (a structure that in gyrencephalic species provides a clear boundary between the claustrum and the neighbouring cortex; for a recent review, see Smith et al. 2018). To overcome the problems that the resulting claustro-cortical continuity has presented, a sustained focus has been on identifying genes that show a differential expression profile in the claustrum relative to surrounding cortical areas. To this end, considerable progress has been made (Mathur, 2014; Mathur et al., 2009; Wang et al., 2017; Watakabe, 2017). Crystallin mu (Crym) expression, for instance, is densely expressed in the insular cortex yet is highly attenuated in the claustrum. Indeed, Crym expression was fundamental to establishing that the claustrum is surrounded on all sides by cortex rather than being juxtaposed with the external capsule (Mathur et al., 2009), as was thought previously. In the same study, the nuclear boundary of the claustrum at the level of the striatum was defined using the expression profiles of parvalbumin, cytochrome oxidase, and the guanine nucleotide binding protein (G protein), gamma 2 (Gng2; Mathur et al., 2009). More recently, Wang et al (2017) compiled a list of 49 genes that are differentially expressed in the mouse claustrum.

Alongside this progression, however, attempts to resolve the issue of whether, or not, the rostral boundary of the claustrum extends beyond the anterior aspect of the striatum have seen limited progress. In the seminal work of Mathur et al. (2009), the apparent absence of parvalbumin and Gng2 expression within the atlas-defined boundary of the rostral claustrum prompted a reassessment of the anatomical boundary of the claustrum to one that did not extend beyond the anterior horn of the neostriatum. Subsequent anatomical and behavioural studies have, for the most part, conformed to the Gng2-based anatomical definition of Mathur et al. (e.g. Smith and Alloway 2010a). In a recent review, however, Smith and colleagues (2018) highlight the importance of reaching a resolution in future studies. Indeed, in another recent review (Dillingham et al., 2017), using a freely available nucleotide sequence expression mouse brain database (Allen Mouse Brain Atlas; available at: http://mouse.brain-map.org/), the expression of a number of genes that were identified as having differential expression in the claustrum (Mathur et al., 2009; Wang et al., 2017) were assessed. Of 49 genes, the striatal - claustrum boundary, delineated either by attenuated expression (e.g. Slit-1, Crym), or enriched expression (e.g. Gng2, Gnb4, latexin), was found to extend considerably rostral to the striatum. Significantly, however, unlike atlas-based delineations, the oval cross section of the claustrum is situated at the ventrolateral aspect of the forceps minor of the corpus callosum, i.e. maintaining its locus in Layer 6 of the insular cortex. Given the multimodal nature of the claustrum (Remedios et al., 2010) and the likelihood that the separate ‘puddles’ of (presumably functional) connectivity act in concert (Smythies et al., 2014), it is all the more important that a consensus in the field relating to its anatomical boundaries is reached.

In the present study, a combination of immunohistochemistry, immunofluorescence, and neuronal tracing was employed to examine the expression patterns of several genes that have been identified as verified claustral markers. One, crystallin mu (Crym), exhibits an attenuated expression in the claustrum relative to surrounding cortex (Mathur et al., 2009), while Gng2 (Mathur et al., 2009) and parvalbumin (Druga et al.; Hinova-Palova et al., 2014a; Mathur et al., 2009; Pirone et al., 2015; Rahman and Baizer, 2007) show enriched expression in the claustrum. In combination with anterograde and retrograde anatomical tracing, the expression profiles of these genes have been reassessed in the rat brain with a focus on establishing the rostral boundary of the claustrum.

## Methods

### Subjects

A total of 41 male Lister Hooded rats (Envigo, UK) with pre-procedural weights of between 230-350g were used in the study. In 6 animals, retrograde neuronal tracer injections targeted nucleus reuniens (RE) and/or the rhomboid (Rh) nucleus of the midline thalamus and in one of these cases, a further retrograde tracer injection targeted the mediodorsal (MD), centromedial (CM) and paraventricular thalamic nuclei (some of these cases used in Mathiasen et al., 2019; **Table 1**). In 2 of the animals with RE/Rh injections the injection site also included a portion of the centromedial thalamic nucleus. Further, in 18 animals, retrograde (n = 13) or anterograde (n = 5) tracer injections targeted the retrosplenial (RSC) or anterior cingulate (Cg) cortices (**Table 1**). In 6 of the animals with tracer injections we further stained for parvalbumin immunofluorescence marker, while a further 16 animals were used for immunohistochemistry (no tracer injections; **Table 1**)

**Table 1.** An overview of the cases used in the study, including details of the type, and target of cortical and thalamic neuronal tracer injections.

| Cases | Injection sites | Tracer | Immunofluorescence |
| --- | --- | --- | --- |
| **Thalamic injections (retrograde tracers)** |  |  |  |
| 215#4 | MD, PV, CM<br>RE/Rh | CtB<br>FB |  |
| 208#9 | RE | FB |  |
| 207#2 | RE/Rh (SMT, IAM) | FB | Parvalbumin, |
| 207#4 | RE/Rh (CM, IMD) | FB |  |
| 207#7 | RE (PVT, CM, IAM) | FB |  |
| 209#10 | RE/posterior<br>hypothalamus | CtB |  |
| 207#1 | RE | FB | Parvalbumin |
| **Cortical injections (retrograde tracers)** |  |  |  |
| 199#29 | RSC | FB |  |
| 225#4 | RSC | CtB |  |
| 225#12 | RSC | CtB |  |
| 223#1 | RSC | CtB |  |
| 198#12 | RSC | FB |  |
| 199#29 | RSC | FB |  |
| 225#1 | RSC | CtB | Parvalbumin |
| 225#1 | Cg | FB | Parvalbumin |
| 222#10 | RSC | CtB |  |
| 222#10 | Cg | FB |  |
| 223#5 | Cg | FB |  |
| 223#26 | Cg | CtB |  |
| FGrsc#1 | RSC | FG | Parvalbumin |
| FGrsc#2 | RSC | FG | Parvalbumin |
| LK#1 | RSC | CtB | Parvalbumin |
| **Cortical injections (anterograde tracers)** |  |  |  |
| 224#30 | RSC | pAAV-CaMKIIa-EGFP |  |
| 224#29 | RSC | pAAV-CaMKIIa-EGFP |  |
| 219#3 | Cg | pAAV-CaMKIIa-hM4D(Gi)-Mcherry | Parvalbumin |
| 219#17 | Cg | pAAV-CaMKIIa-hM4D(Gi)-Mcherry |  |
| 219#21 | Cg | pAAV-CaMKIIa-hM4D(Gi)-Mcherry |  |
| **Immunohistochemistry** | **Marker** | **Plane** |  |
| HPCLA1 | Crym, Gng2 | Coronal |  |
| HPCLA2 | Crym, Gng2 | Coronal |  |
| RCLA1 | Crym, Gng2 | Coronal |  |
| RCLA2 | Gng2 | Coronal |  |
| RCLA3 | Gng2, Crym | Coronal |  |
| RCLA4 | Gng2 | Coronal |  |
| RCLA5 |  | Coronal |  |
| SHPC1 | PV | Coronal |  |
| BiFlex1 | PV, Crym | Coronal |  |
| BiRCLA1 | PV, Crym | Coronal |  |
| BiRCLA2 | PV | Coronal |  |
| BiRCLA3 | PV, Crym | Coronal |  |
| CLArsc2 | PV | Horizontal (axial) |  |
| WK2 | PV | Horizontal (axial) |  |
| LH1 | Gng2 | Coronal |  |
| LH2 | Gng2 | Coronal |  |

### Compliance with Ethical Standards

Animal husbandry and experimental procedures were carried out in accordance with the European Community directive, 86/609/EC, and the Cruelty to Animals Act, 1876, and were approved by the Comparative Medicine/Bioresources Ethics Committee, Trinity College, Dublin, Ireland, and followed LAST Ireland and international guidelines of good practice or, for those experiments that were performed at Cardiff University, in accordance with the UK Animals (Scientific Procedures) Act, 1986 and associated guidelines, the EU directive 2010/63/EU, as well as the Cardiff University Biological Standards Committee.

### Surgical procedures

Anaesthesia was induced and maintained with isoflurane (5% and 1–2%, respectively) combined with oxygen (2 L/minute). Animals were then placed in a stereotaxic frame (Kopf, Tujunga, CA, USA) and chloramphenicol eye ointment (Martindale Pharmaceuticals, Romford, UK) was topically applied to the eyes to protect the cornea. Pre-surgical analgesia (Metacam, 1 mg/kg; Boehringer Ingelheim, Germany) and antibiotics (Enrocare; Animal Care Ltd., York, UK) were administered subcutaneously. The scalp was incised and cleaned before crainotomies were made, large enough to permit advancement of a Hamilton syringe into thalamic or neocortical regions. For retrograde tracing we injected either Fast Blue (FB; Polysciences), Fluorogold (FG; Santa Cruz Biotechnology, Heidelberg, Germany), or cholera-toxin b (CtB; List Biological Labs Ltd., California, USA). For anterograde tracing we injected an adeno-associated virus expressing either MCherry (pAAV-CaMKIIa-hM4D(Gi)-Mcherry) (AAV5), or GFP (pAAV-CaMKIIa-EGFP; AAV5) as a fluorescent marker (Addgene, Cambridge Massachusetts, US).

Following tracer-specific survival times, ranging from 5 days to 7 weeks (the latter for viral injections in anterior cingulate only), rats were deeply anaesthetised with sodium pentobarbital (Euthanimal) and perfused transcardially with ice cold 0.1M phosphate buffered saline (PBS) followed by either 2.5% or 4% paraformaldehyde in 0.1M PBS.

### Immunohistochemistry (IHC)

In cases used for immunohistochemistry (with no tracer injections), the brains were removed and post-fixed for 48 hours before being transferred to a 25% sucrose in 0.1M PBS solution for 1-2 days for cryoprotection. Sections of 40μm were cut on a cryostat (Leica CM1850) with one 1-in-4 series mounted directly on to double gelatine-subbed microscope slides. Of the remaining 3, 1-in-4 series, one or more of: an anti-Gng2 polyclonal antibody raised in rabbit (Sigma-Aldrich Ireland Ltd; Wicklow, Ireland), an anti-Crym monoclonal antibody raised in mouse (Novus Biologicals; Abingdon, UK), and an anti-parvalbumin monoclonal raised in mouse (Swant Inc. Marly, Switzerland) were used to immunolocalise the respective proteins.

Initially, endogenous peroxidases were removed from free-floating brain sections through reaction in a quench solution containing 10% methanol and 0.3% hydrogen peroxide in distilled water. Following washes in PBS and subsequently PBST (0.05% Triton X-100 in 0.1M PBS), the sections were agitated in a 4% solution of normal horse serum in 0.1M PBS for 2 hours. Sections were then transferred to a 1:200 dilution of either anti-Crym or anti-Gng2 in 0.1M PBST with 1% normal horse serum and agitated at 4°C overnight. Following washes in PBST, sections were transferred to a 1:250 dilution of biotinylated horse-anti-mouse IgG (for sections reacted against Crym; Vector Labs, Peterborough, UK) or biotinylated horse-anti-rabbit IgG (for sections reacted against Gng2; Vector Labs, UK) for 2 hours. Sections were then washed in PBST before undergoing signal amplification through incubation in the Vectastain ABC solution (Vector Labs, Peterborough, UK) for 2 hours. Following washes in PBST and subsequently PBS, sections were agitated overnight at 4°C. Immunoreactivity was visualised using the chromagen diamino benzidine (DAB; Vector Labs, Peterborough, UK) and in some cases, the signal was intensified with by adding nickel chloride to the DAB solution. Sections were then washed in PBS, mounted, and left to dry at room temperature before being dehydrated in ascending alcohols, cleared in xylene, and coverslipped with DPX mounting medium (Sigma-Aldrich, Gillingham, UK).

### Immunofluorescence staining

In cases with anterograde or retrograde tracer injections, brains were post-fixed for 4 hours before being transferred to a 25% sucrose solution overnight. Sections of either 40μm or 50μm were cut in the coronal place with a freezing microtome with one 1:4 series used for Nissl staining, a second series used for visualization of the tracers and, in some cases, remaining series were used for further immunofluorescence staining (**Table 1**).

For visualization of neuronal tracers, brain sections were washed in PBS and PBS-TX followed by incubation with the relevant primary antibody (rabbit anti choleratoxin (Sigma-Aldrich UK) or rabbit anti-Mcherry (1:2000 dilution; Abcam, Cambridge, UK) overnight. Following PBS-TX washes the sections were incubated with the secondary antibody (1:200 dilution; goat anti-rabbit DyLight 594; Vector Laboratories; Peterborough, UK), washed in PBS, mounted and coverslipped either directly with Fluoromount (Sigma-Aldrich, Gillingham, UK) or alternatively, following dehydration in ascending alcohols, with DPX mounting medium (ThermoFisher, Waltham, MA).

In a number of these cases (see **Table 1**) sections were further stained for mouse anti-parvalbumin, (1:10000 dilution; Sigma-Aldrich, UK) in a 1% NGS (company) PBS-TX solution following 90 min in a 5% NGS solution. Sections were incubated overnight, washed in PBS-TX and incubated with the relevant secondary antibody (goat anti-mouse A488 (Abcam, Cambridge, UK) in a 1% NGS 1:200 PBS-TX concentration.

### Microscopy and imaging

Brain sections were imaged in brightfield at 20x magnification using a Leica slide scanner (Aperio AT2), visualised in Aperio ImageScope (version 12.3.2.8013). For fluorescence microscopy two systems were used. Either a Leica DM6000 B microscope with an attached Leica DFC350 FX digital camera with acquisition software (LAS AF image, Leica), or a Leica DM5000 B microscope with an attached Leica DFC310 FX digital camera. Images were adjusted for brightness and contrast in Corel Photo Paint X5 or FIJI (‘*fiji is just imageJ*’ freely available software available from https://imagej.net/Fiji/Downloads). Pixel density heat maps were generated in Fiji; images were converted to 8-bit and median filtered before applying a 16-colour LUT.

#### Nomenclature

Based on their recent guidance and clarification on the issue of how to consider the claustrum in relation to the endopiriform nuclei, we follow the classification of Smith and colleagues (2018) and consider a claustrum-endopiriform complex comprising the claustrum proper and the dorsal endopiriform nucleus (DEn), with the claustrum proper comprising dorsal (dCLA) and ventral (vCLA) subdivisions. In addition to atlas-based (Paxinos and Watson, 2005) definitions of the insular/orbital region, Nissl stained sections were used to delineate the border between the insular and orbital cortices. The lateral orbital cortex display prominent cytoarchitectonical differences from the insular cortex, such as a more densely packed layer 5 and a less sharp border between layers 2 and 3 (Van De Werd and Uylings, 2008).

## Results

### Anatomical boundary - Parvalbumin (PV) (IHC)

#### Coronal plane

Neuropillar parvalbumin expression in the agranular insular cortex is characterized by attenuated expression with the exception of a densely-labeled fiber plexus in layer 5 (**Fig. 1A-D**). Contrasting dense expression of parvalbumin was present in the neuropil of the vCLA, distinctly revealing its core (Smith et al., 2018). Parvalbumin-immunoreactive neuron density was found to be sporadic but uniformly distributed across the insular cortex and vCLA with no discernible inter-laminar difference (**Fig. 1C-D**).

**Figure 1.**
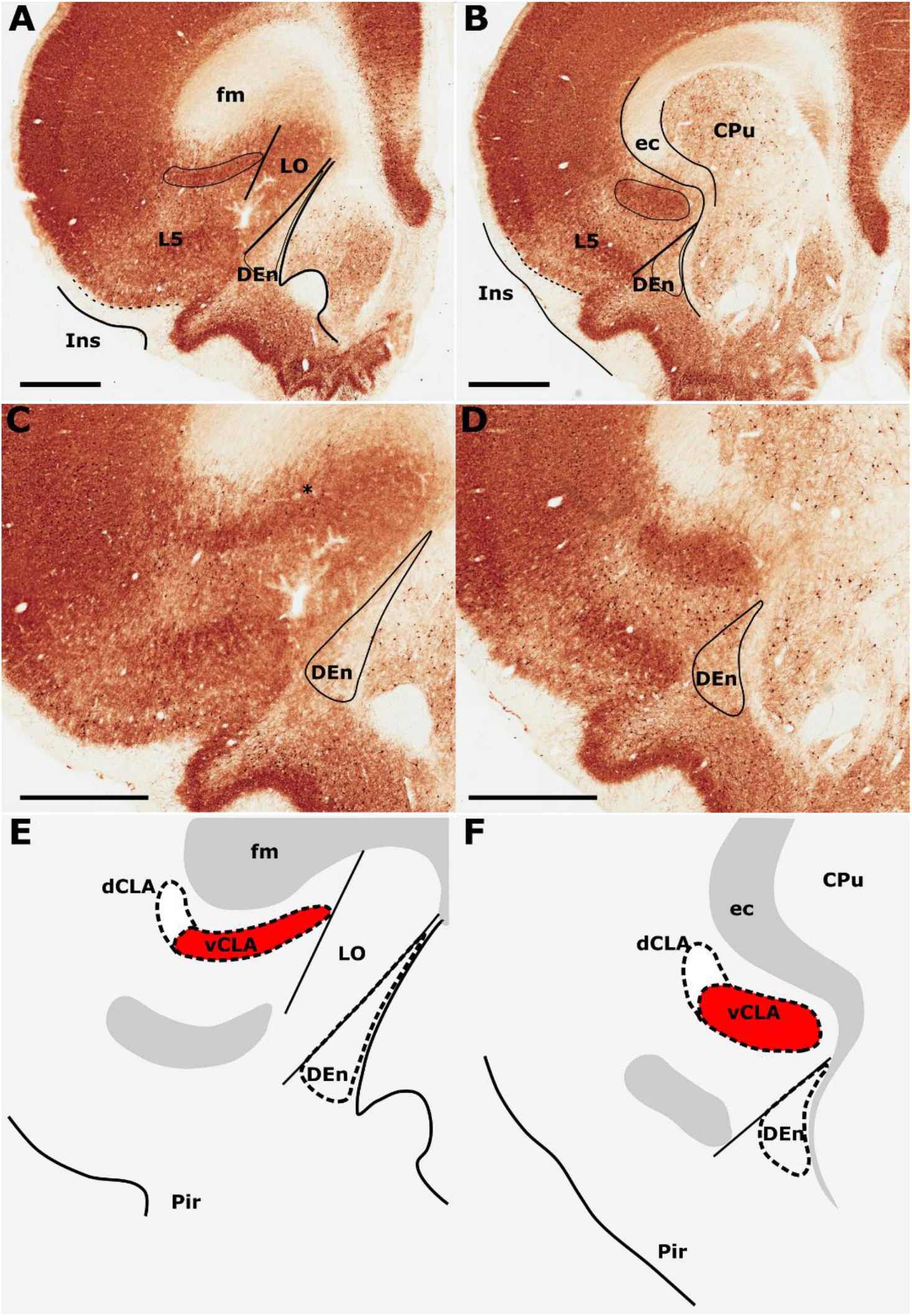
Differential parvalbumin expression in the insular region aides the delineation of the anatomical border of the claustrum. **A-D** show low (**A, B**) and high (**C, D**) magnification photomicrographs of parvalbumin immunoreactivity in brain sections rostral to the striatum (**A, C**) and at the level of the striatum (**B, D**). At the level of the striatum, dense expression of parvalbuminergic neuropil in the ventral claustrum (vCLA) contrasts with weaker expression in the neighbouring layer 6 of the insular cortex (B, D). Rostral to the striatum, dense parvalbuminergic neuropil is again observed deep to layer 6 of the insular cortex (Ins). Parvalbumin is expressed in the vCLA but does not allow for the delineation of the borders of the dorsal endopiriform nucleus (DEn), or the dorsal claustrum (dCLA) whose boundaries are estimated in **C-D**. Images **E-F** are schematic representations of **C-D**, respectively. Red fill represents the part of the complex that can be delineated using parvalbumin immunolocalization. Note in **A** and **C** that due to dense expression in the lateral orbital cortex (LO), the border between the medial extent of vCLA and LO is not easily determinable (asterisk). Abbreviations: CPu, caudate/putamen; ec, external capsule; fm, forceps minor of the corpus callosum; Pir, piriform cortex. Scale bars = 1000 μm.

The insular cortex is bordered caudally by the peri- and ectorhinal cortices, while the rostral boundary of the insular cortex interfaces with the orbital cortices (**Fig. 1A, C**). Both orbital and rhinal cortices regions exhibit a uniformly higher density of parvalbumin neuropillar immunoreactivity across layers 4-6, albeit again with increased expression in layer 5. The transition of insular to peri/ectorhinal cortex matches closely with the caudal apex of the vCLA (**Fig. 2**), i.e. a continuous extension of claustral PV expression into the rhinal cortices was not present.

**Figure 2.**
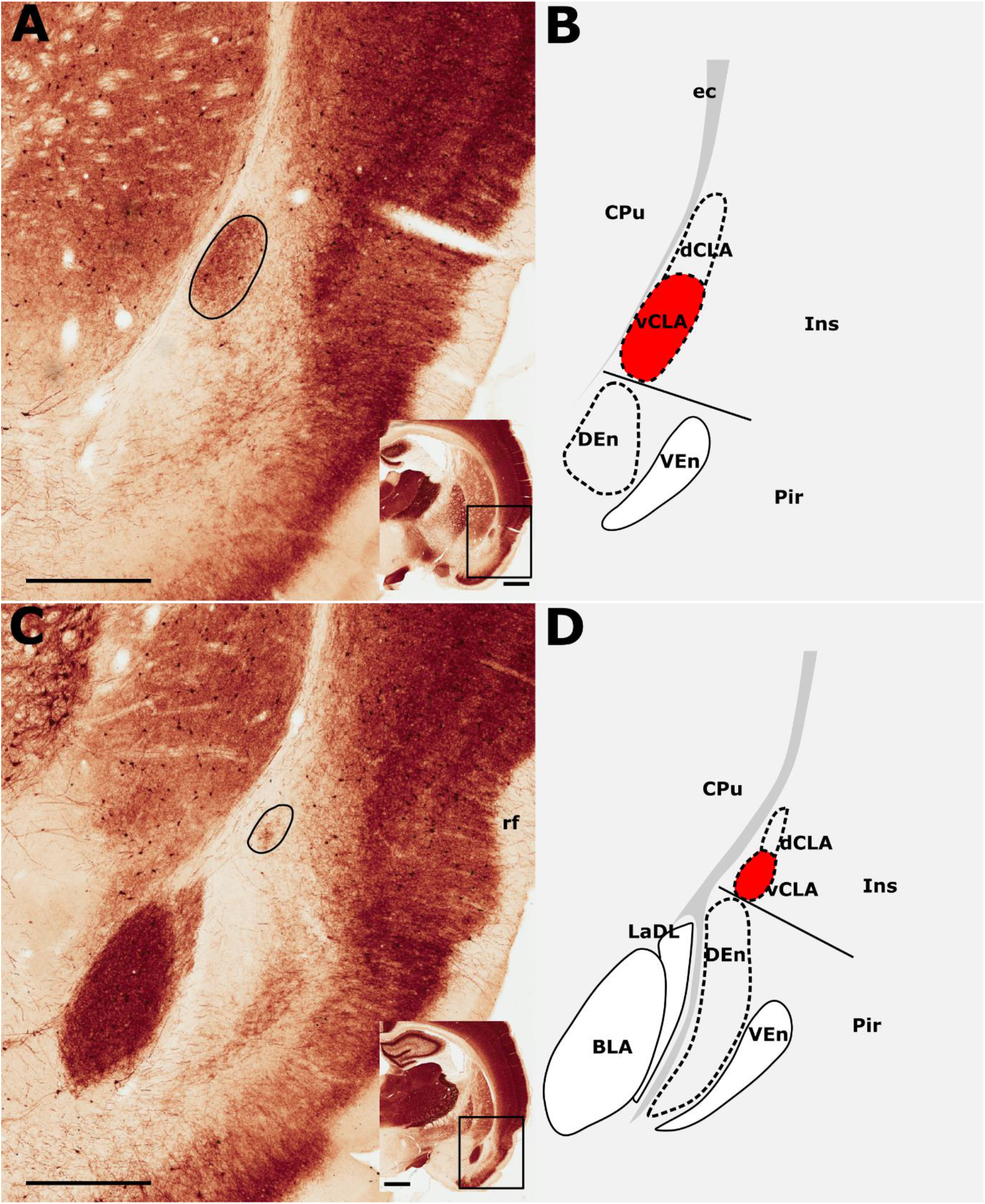
Parvalbumin expression showing the caudal extent of the ventral claustrum (vCLA). **A**, At a level approximately −1.8 mm from bregma (see low magnification inset), the claustrum is situated within layer 6 of the insular cortex (Ins). Note the absence of label in the dorsal claustrum (dCLA). **B**, Further caudally, at the level to the anterior amygdala complex (approximately −2.5 mm from bregma; see low magnification inset), the claustrum is significantly reduced in cross sectional area. The vCLA was not present caudal to this coronal section. Schematic representations of **A** and **C** are shown in **B** and **D**, respectively, with red fills representing parvalbumin expression in the claustrum. Abbreviations: BLA, basolateral amygdala nucleus; CPu, caudate putamen; DEn, dorsal endopiriform nucleus; ec, external capsule; Ins, insular cortex; LaDL, lateral amygdaloid nucleus, dorsolateral part; Pir, piriform cortex; VEn, ventral endopiriform nucleus. Scale bars in A, C = 500 μm; insets = 1000 μm.

At the anterior horn of the neostriatum, parvalbumin expression in vCLA remained dense with no apparent reduction in cross-sectional area. At this coronal level (approximately represented by the +2.52 (from Bregma) plate in Paxinos and Watson, 2005), the ovoid cross-section of the claustrum is more horizontally oriented (**Fig. 3B-C**), being elongated within the arch of the external capsule. Immediately rostral to the anterior horn of the neostriatum, the putative vCLA was still present, maintaining its position beneath the forceps minor of the corpus callosum (within layer 6 of the insular cortex). Further rostrally, the lateral orbital cortex emerges to laterally displace both the insular cortex and the vCLA to a position progressively more ventrolateral with respect to the forceps minor (**Fig 1A, C**).

**Figure 3.**
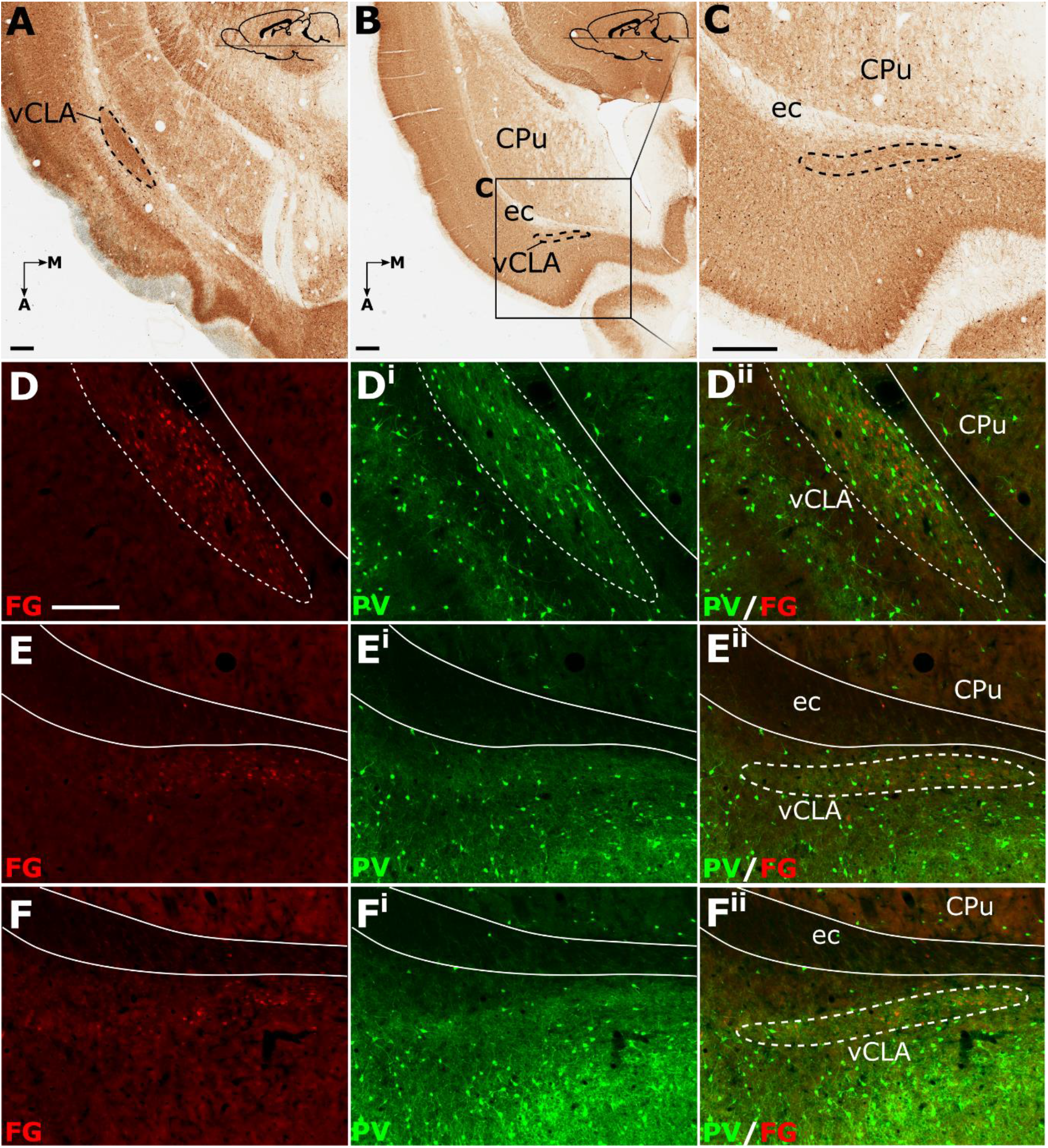
Parvalbumin expression in horizontal sections (**A-D**) provided complementary findings to those derived from coronal sections. Unlike coronal or sagittal series, the horizontal plane allows for visualization of much of the anterior-posterior extent of the ventral claustrum (vCLA) in a single section (**A**). Low magnification (**B**) and high magnification (**C**) images of the rostral extent of the vCLA (dashed line) in a comparatively more dorsal section than **A**. Dual fluorescence, combining parvalbumin expression (green) and retrograde labelling following injections of Fluoro-gold into the retrosplenial cortex (red), showed that, at striatal levels (**D-D^ii^**, corresponding to that in **A**), dense retrograde label overlaid claustral parvalbumin enrichment. Rostral to the striatum (**E-E^ii^** and **F-F^ii^**) retrograde labelled cell soma again overlaid parvalbumin enrichment corresponding to the claustrum, albeit with fewer retrogradely labelled soma and weaker parvalbumin expression. Scale bar in A-C = 500 μm. Scale bar in **D** (applies to all fluorescent images) = 250 μm.

In our analyses of parvalbumin across multiple coronal cases, the vCLA was consistently found to extend approximately 500 μm anterior to the anterior horn of the striatum. It is worthy of note that parvalbumin expression in the lateral orbital cortex was uniformly dense across its layers, such that the medial extent of the vCLA and the lateral orbital cortex appeared continuous; it was therefore difficult to determine this border between regions (**Fig. 1A, C**).

#### Horizontal (Axial) plane

Relative to the midline, the position of the claustrum courses approximately 2 mm medially from its caudal position at bregma to its rostral position at the anterior horn of the neostriatum (Paxinos and Watson, 2005), such that visualisation of the nucleus in the true-sagittal plane is only moderately beneficial in examining its continuity in the rostro-caudal axis. The dorsal-ventral position of claustrum, however, remains relatively consistent along this rostro-caudal extent such that visualisation of large portions of its continuity in the same plane of section is possible (e.g. 2 mm in **Fig. 3A**). Retrograde tracer injections (Fluoro-gold) into the retrosplenial cortex combined with parvalbumin immunofluorescence in horizontal brain sections to further assess the rostral extent of vCLA. At striatal levels both parvalbumin expression and distributions of retrograde cell soma label clearly demarcated the claustral area. Beyond the anterior horn of the neostriatum, the vCLA arches upwards beneath the forceps minor. As a result, retrograde label and parvalbumin expression were observed in comparatively dorsal horizontal sections (**Fig. 3B-C**). At these dorsal levels, unlike in coronal sections in which contrast is present between dense claustral parvalbumin expression and weak expression in the immediately adjacent layer 6 of the insular cortex, the claustrum in horizontal sections is bordered by comparably dense cortical parvalbumin expression. At both striatal levels (**Fig. 3D-D^ii^**), and rostral to the striatum (**Fig. 3E-E^ii^** and **F-F^ii^**), retrogradely labelled cell bodies were present in the claustrum and in distributions that closely matched claustral parvalbumin expression.

### Anatomical boundary - Crystallin mu (Crym) (IHC)

In findings that are consistent with reports in the mouse (Wang et al., 2017) and rat (Mathur et al., 2009), expression of Crym was dense in the insular cortex at striatal levels of the telencephalon but markedly reduced in the vCLA (**Fig. 4C, G; Fig. 5**). In the putative dCLA, Crym-immunoreactive neuropil was reduced but to a lesser degree, while DEn was not discernible as the intensity of Crym immunoreactivity was similar to that in the neighbouring piriform cortex (**Fig. 5A-C**). Within the insular region, particularly high densities of Crym-immunoreactive cell bodies and neuropil were distributed around the circumference of the vCLA/dCLA complex. Within the core of the vCLA and dCLA, the distribution density of Crym-immunoreactive cell bodies was considerably reduced with just a few scattered ectopic Crym-positive soma (**Fig. 4E-H**; **Fig. 5**), although rostral to the striatum, the density of these ‘ectopic’ cortical soma was higher (**Fig. 4A-D**).

**Figure 4.**
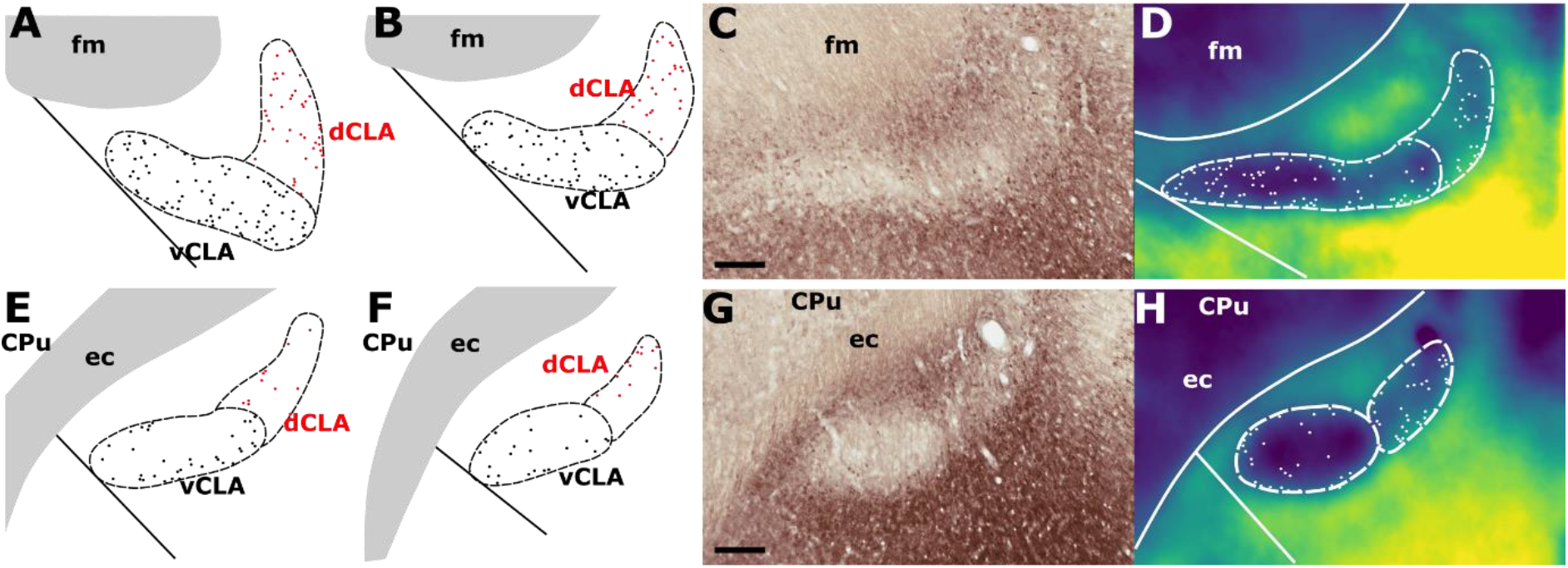
**A-B** Schematic representations of crystallin mu (Crym) delineated boundaries of the claustrum anterior to the striatum (CPu), highlighting the density of ectopic Crym-positive cell soma within the core of the ventral (vCLA; black) and dorsal claustrum (dCLA; red). **C** shows representative Crym staining in the claustrum/insular while **D** shows a pixel density heat map of C highlighting 1. The difficulty associated with determining the boundary between vCLA and dCLA; and 2. Cortical fibers crossing the claustrum to join the internal capsule. **E-H** show equivalent panels from striatal levels in which the number of ectopic Crym-positive cell soma is reduced and the boundary between vCLA and dCLA is more distinct. Abbreviations: ec, external capsule; fm, forceps minor of the corpus callosum. Scale bars = 200 μm.

**Figure 5.**
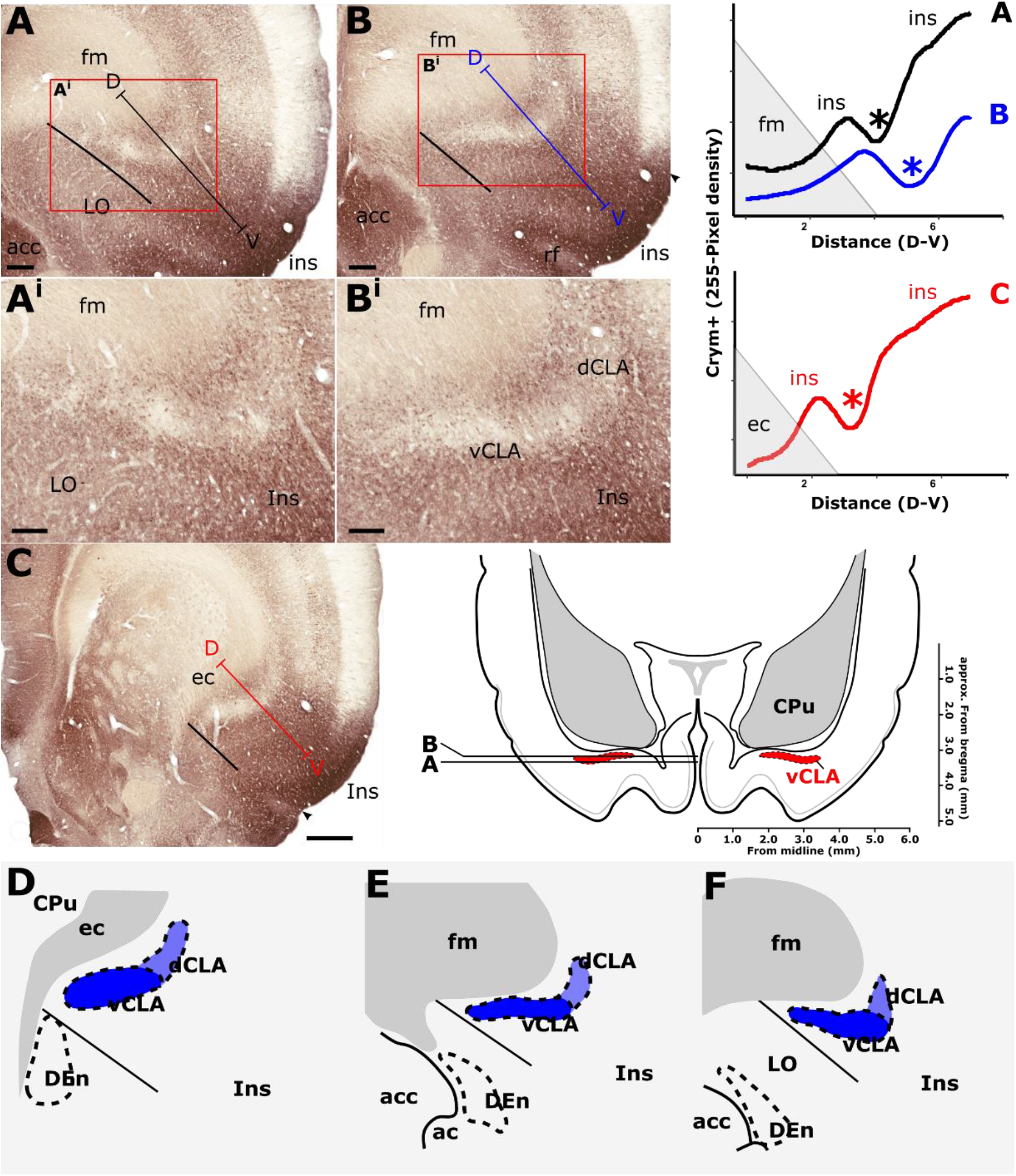
Crystallin mu (Crym) is a cortical marker that is expressed particularly strongly in the insular cortex. In contrast, expression in the claustrum is considerably reduced providing contrast for delineation of the anatomical boundary of the claustrum. Pixel density plots through the insular cortex at both striatal levels (C; red), and rostral to the striatum (**A, B**; black and blue, respectively) show cortical peaks either side of a claustral trough (asterisks). Central schematic diagram shows approximate coronal levels of photomicrographs in **A-B. D-F**: Schematic representations of Crym-based delineation of the claustrum-endopiriform complex. Delineation of the dorsal endopiriform nucleus (DEn) is not possible using Crym, however, vCLA, and to a lesser degree dCLA are (See also **Fig. 3**). Abbreviations: acc, nucleus accumbens; CPu, caudate/putamen; ec, external capsule; fm, forceps minor of the corpus callosum; Ins, insular cortex; LO, lateral orbital cortex. Scale bars in A, B = 300 μm; A^i^, B^i^ = 200 μm; C = 600 μm.

Consistent with past dual-immunofluorescence (Crym and parvalbumin) experiments (Mathur et al., 2009), differential expression of Crym in vCLA relative to the surrounding insular cortex delineated an anatomical boundary that closely matched that derived from our parvalbumin expression profile (**Fig. 5C-E**; supplementary **Fig. S1**), forming an increasingly elongated ovoid cross-section in the coronal plane towards the anterior horn of the neostriatum. Beyond the striatum, the Crym-based vCLA boundary formed a horizontally oriented ovoid beneath the forceps minor of the corpus callosum while further rostrally it was found to apex ventrolaterally beneath the forceps minor (while remaining confined to the boundary of the insular cortex; **Fig. 5D-E**). Unlike the parvalbumin expression profile, however, the Crym profile enabled a clear delineation of the boundary between the vCLA (weak Crym expression) and the lateral orbital cortex (dense Crym expression), with the finding that vCLA did not extend into the lateral orbital cortex but remained confined to the boundaries of the insular region (**Fig. 5A-B**). Crym expression was also found to be reduced in dCLA (**Fig. 5**), which meant that the precise vCLA-dCLA transitional boundary was not clear; an issue that was also contributed to by the presence of Crym-immunoreactive fibers ascending to the internal capsule (Coizet et al., 2017; **Fig. 5A^i^-B^i^**).

### Anatomical boundary - Gng2 (IHC)

Neuropillar Gng2 immunoreactivity was found to be densely distributed throughout the insular cortex (**Fig. 6A-D**). The densest Gng2-immunoreactivity was present in the superficial-most layers and was reduced in layers 5 and 6 which contrasted with the dense vCLA immunoreactivity (**Fig. 6A-D**). Dense expression was observed in layer 2 of the piriform cortex but weak expression in layer 3 again provided contrast with denser Gng2 immunoreactivity in DEn (**Fig. 6A-D**).

**Figure 6.**
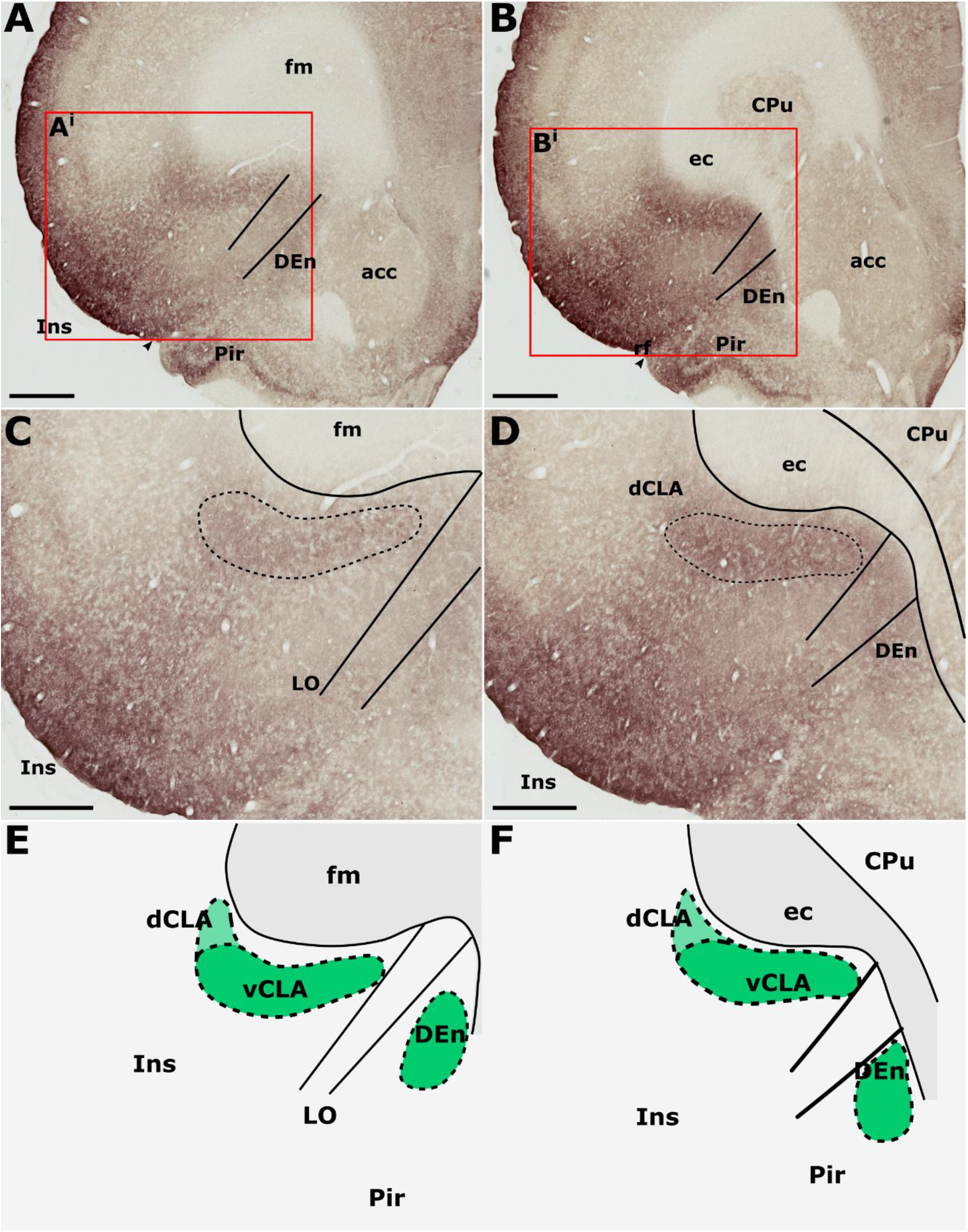
Guanine nucleotide binding protein (G protein), gamma 2 (Gng2), is expressed throughout the insular cortex but relatively weakly in layers 5/6. At striatal levels (**B, D**), dense expression is observed in the ventral claustrum (vCLA) and dorsal endopiriform nucleus (DEn). Expression in the dorsal claustrum (dCLA) but it is relatively weaker (**D**). The same distribution of expression is evident anterior to the striatum (**A, C**) albeit with weaker expression throughout the claustrum-endopiriform complex. Note in A and C, the separation of DEn from vCLA with the emergence of the lateral orbital cortex (LO). **E-F** are schematic representations of Gng2 expression (green) rostral to the striatum and at striatal levels, respectively. Abbreviations: acc, nucleus accumbens; CPu, caudate/putamen; ec, external capsule; fm, forceps minor of the corpus callosum; Ins, insular cortex; Pir, piriform cortex. Scale bars in A, B = 800 μm; C, D = 500 μm.

Gng2 immunoreactivity delineated a vCLA boundary that was consistent with both parvalbumin and Crym, albeit with a less-well pronounced margin (**Fig. 6E-F**). Indeed, manual registration of serial sections that had been immunohistochemically (DAB) reacted for either Crym or Gng2, revealed expression profiles of vCLA and dCLA Gng2 enrichment that closely matched (at all claustral levels) the region of Crym attenuation in the corresponding section (see supplementary **Fig. S2**). At striatal levels, Gng2 enrichment in DEn was continuous ventrally with vCLA although with denser expression in vCLA (**Fig. 6B, D**), so that the boundary between the two nuclei at the piriform/insular boundary was distinct. Rostral to the anterior horn of the striatum, DEn was no longer continuous with vCLA, and the two regions became progressively separated by the emergence of the lateral orbital cortices, i.e. the extent of vCLA and DEn remained confined to insular and piriform cortices, respectively (**Fig. 6A, C, E**). At this anterior-posterior level, Gng2 expression in vCLA formed a horizontally-oriented ovoid beneath the forceps minor of the corpus callosum with the dCLA arching around its ventrolateral border. As with PV and Crym, further rostrally, vCLA became more restricted in cross-sectional area and situated more laterally with respect to the forceps minor.

### Anatomical boundary - Neuronal tracer injections

Pressure injections of either retrograde (FB, CtB or FG) or anterograde (viral) neuronal tracers were made targeting either the retrosplenial or anterior cingulate cortices, revealing a consistently dense pattern of label along the rostro-caudal extent of vCLA (see **Table 1**).

Cases in which multiple FB, FG or CtB injections were made unilaterally along the extent of the retrosplenial cortex or anterior cingulate cortex (more confined injections) resulted in dense retrograde label in the ipsilateral claustrum. Although weak, retrograde label was present in the claustrum of the contralateral hemisphere at both striatal levels, as well as rostral to the striatum (see supplementary **Fig. S3**). The distribution of retrogradely labeled cell bodies was confined to the ventral claustrum, i.e., it did not extend into the dorsal claustrum, or ventrally to the DEn. Significantly, the distribution of retrograde label in vCLA extended beyond the anterior horn of the neostriatum, delineating a boundary consistent with that determined from IHC analyses (**Fig. 7A-A^ii^**). In cases involving retrograde injections targeting the anterior cingulate cortex, dense cell labelling was present in the claustrum between 0.4-0.6 mm rostral to the anterior tip of the striatum. The cell labelling that resulted from injections in the retrosplenial cortex extended to comparable rostral levels, although with varying cell density.

**Figure 7.**
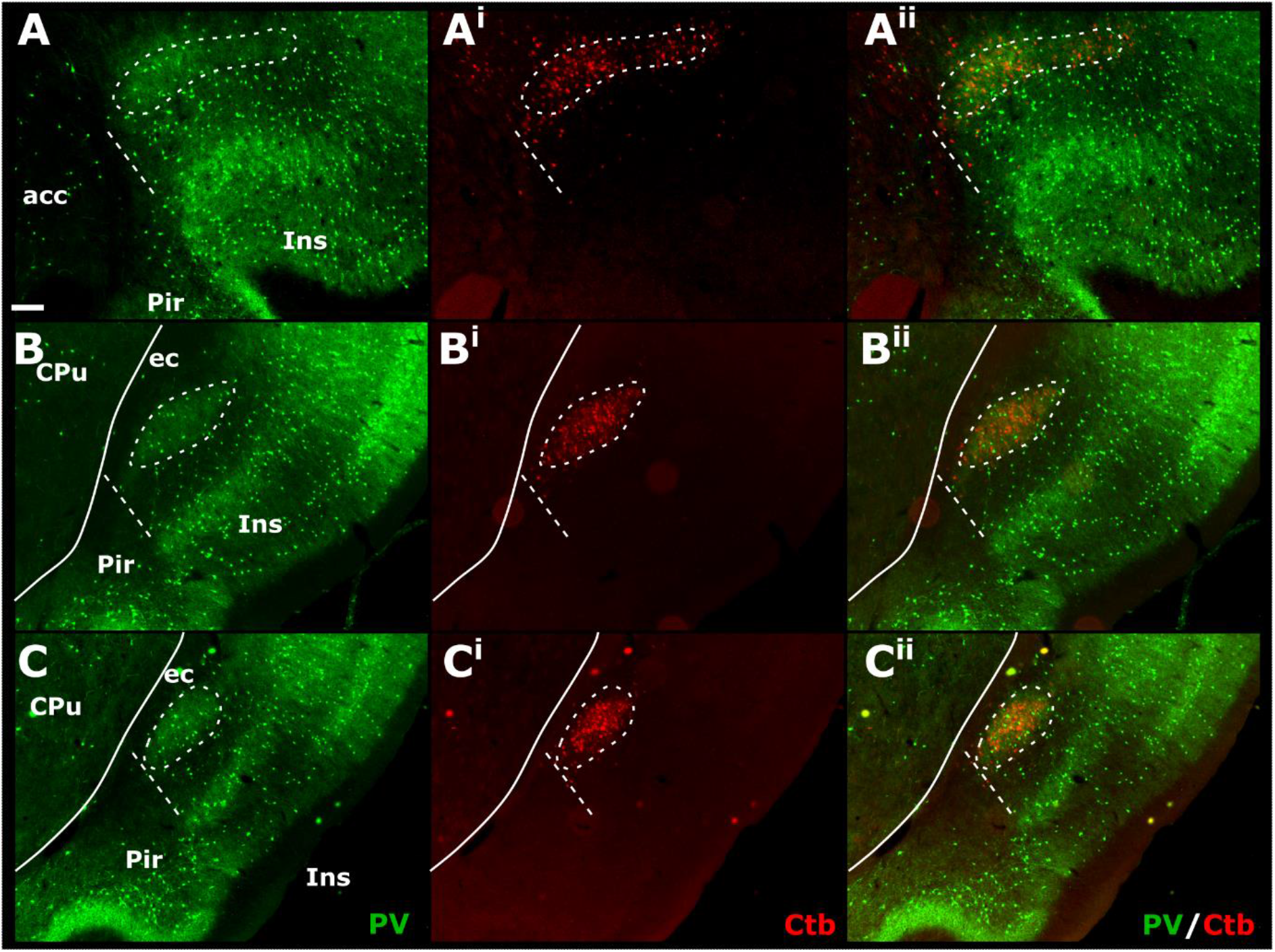
Tracer injections of the non-toxin subunit B of cholera toxin (CtB) within the retrosplenial cortex resulted in dense cell soma label in the ipsilateral claustrum in a distribution that overlapped with claustral parvalbumin expression (for a comparable case using Fluoro-gold, see supplementary **Fig. S4** and for injections sites see supplementary **Fig. S5**). At anterior-posterior (AP) levels rostral to the anterior horn of the neostriatum (CPu, A-A^ii^) parvalbumin and CtB label delineated a claustral border (white dashed line) that was horizontally oriented within the arch of the forceps minor of the corpus callosum (fm). At rostral and mid-striatal AP levels (B-B^ii^ and C-C^ii^, respectively), the claustral border was more vertically oriented alongside the external capsule (ec). Abbreviations: Scale bar = 200 μm.

Injections of anterogradely transported pAAV-CaMKIIa-hM4D(Gi)-Mcherry (serotype 5) confined to the anterior cingulate cortex (bilaterally in three cases; see **Table 1**), resulted in dense fibre/terminal labeling along the rostro-caudal extent of the claustrum (**Fig. 8**), revealing a rostral extension of the claustrum beyond the anterior horn of the neostriatum that closely matched the distribution of retrograde label observed in CtB and FB cases. In these cases, however, the dense “plexus” of fibre label in the claustrum (deep to the insular cortex) was continuous with more widespread, diffuse fibre labelling in the orbitofrontal cortex which, rostral to the level of claustrum, centred in a fibre plexus in the deepest lamina of the lateral orbital cortex together (including more superficial labelling; **Fig. 8A_1_-A_3_** and **B_1_-B_3_**). This orbital portion correspond to what has previously been suggested to constitute the rostral portion of the claustrum (Paxinos and Watson, 2005). In two further cases, injection of pAAV-CaMKIIa-EGFP (serotype 5) into the retrosplenial cortex (unilaterally) resulted in a more restricted fibre distribution, as no dense fibre label was present in the lateral orbital cortex.

**Figure 8.**
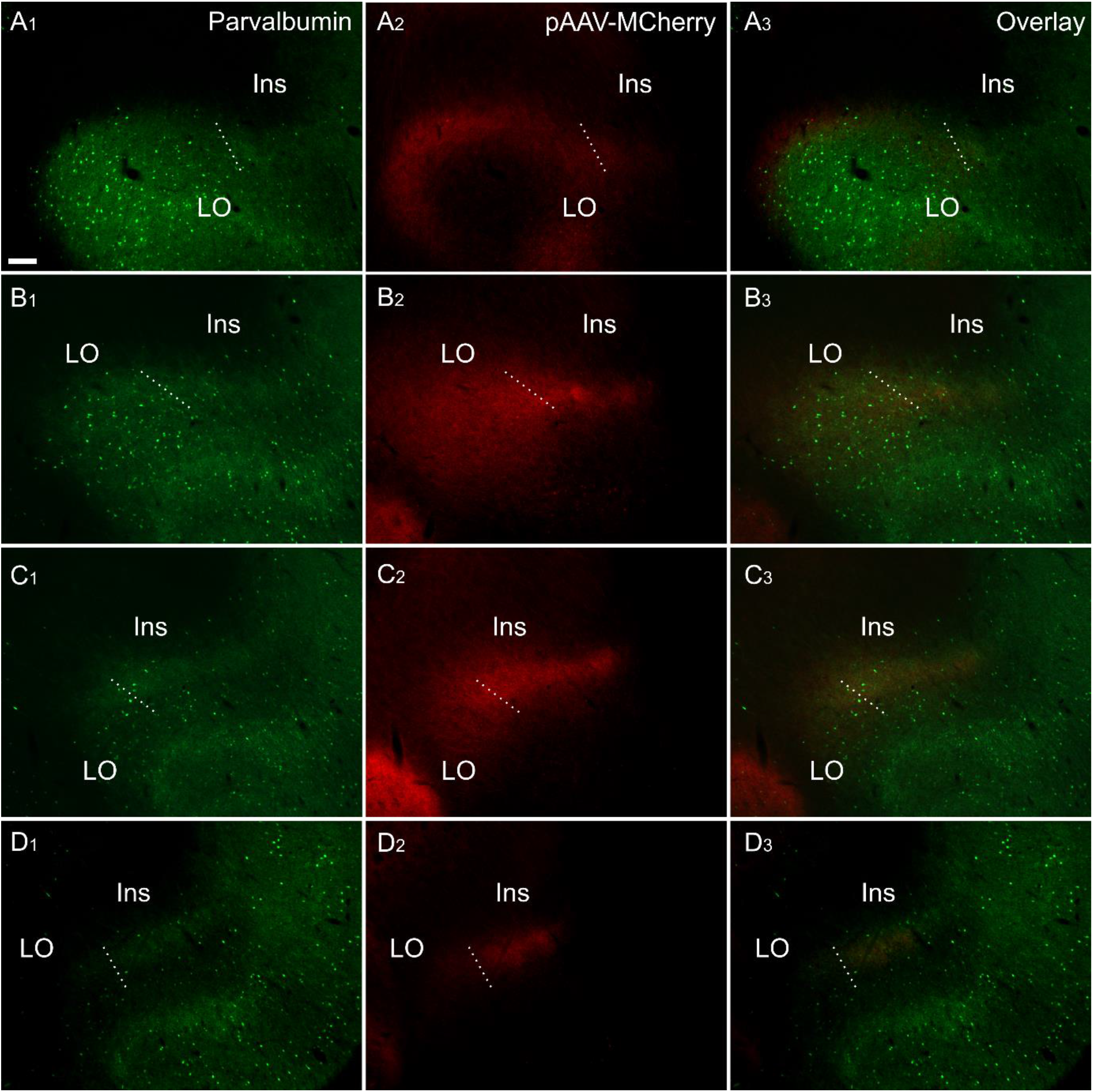
Fibre label resulting from multiple injections (bilateral) of an anterograde viral tracer (pAAV-CaMKIIa-hM4D(Gi)-Mcherry) in the anterior cingulate cortex (case 219#3; column 2) shown in sections colabelled for parvalbumin (column 1). The four rows (**A-D**) show photomicrographs of the claustrum at four anteroposterior levels separated by 200 μm. Row **D** is at the most rostral portion of striatum, Rows **C-A** rostral to striatum. Terminal fibre label co-localized with parvalbumin in a plexus that extends at least two sections rostral to striatum. The dotted line indicates the border between lateral orbital and insular cortices. Abbreviations: Ins, insular cortex; LO, lateral orbital cortex. “pAAV-MCherry” is an abbreviation for pAAV-CaMKIIa-hM4D(Gi)-Mcherry. Scale bars = 200 μm.

In cases in which either retrograde (CtB, FG or FB), or anterograde (pAAV-CaMKIIa-hM4D(Gi)-Mcherry) injections were made into the retrosplenial cortex (CtB) or anterior cingulate cortex (viral tracer and FB), we reacted the sections for parvalbumin (see **Fig. 8** for the anterograde tracing). In these dual-fluorescence cases, immunolocalization distributions of tracer label again closely matched parvalbumin neuropillar immunoreactivity in the vCLA. The distribution of retrograde labelled cell bodies in CtB and FB cases, as well as anterograde fibre/terminal label in DREADDs-mCherry cases, rostral to the anterior horn of the neostriatum, was closely aligned with our parvalbumin-based definition of the rostral claustral area, as described above. Interestingly, the fibre label in the deep layer 6 of the lateral orbital cortex, which resulted from anterograde tracer injections in the anterior cingulate (see above), was shown to a large extent to overlap with a portion *devoid* of parvalbumin neuropillar label (**Fig. 8A_1-3_**).

### Anatomical boundary - thalamocortical connectivity

Retrograde tracer injections (FB or CtB; see **Table 1**), centred in the nucleus reuniens/rhomboid nuclei of the midline thalamus, resulted in dense retrograde label in the insular cortex. A comparable pattern of labelling was seen following an injection centred in the mediodorsal, paraventricular and centromedial thalamic nuclei (**Fig. 9**; see also supplementary **Fig. S5**).

**Figure 9.**
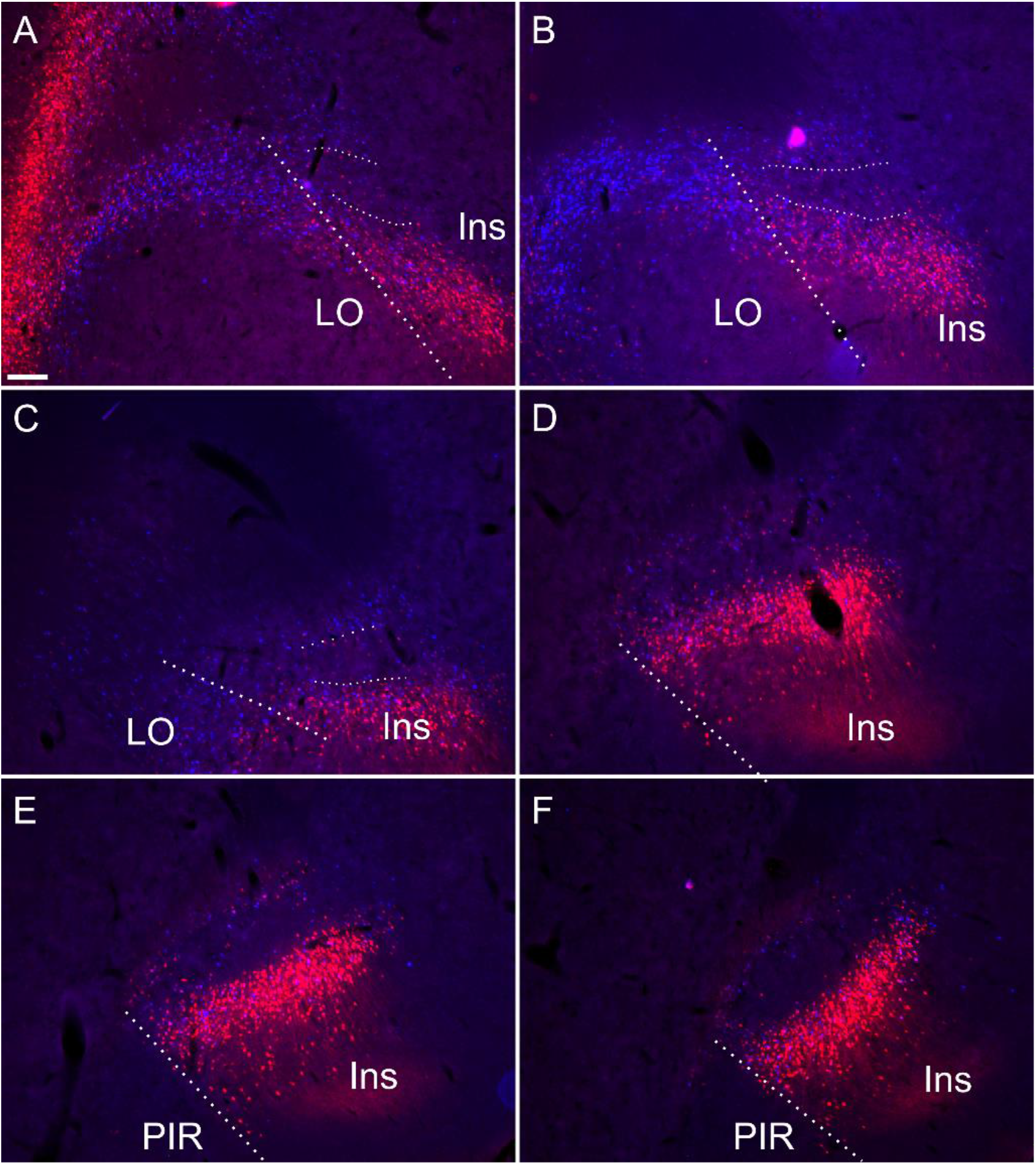
Retrograde soma label, in the insular and orbital cortices, resulting from cholera-toxin b (CtB, red) and Fast Blue (FB, blue) injections centred in mediodorsal/paraventricular/centromedial thalamic nuclei (Ctb) and nucleus reuniens/rhomboid (blue; see supplementary **Fig. S5** for injection sites corresponding to this case). The dense soma label encapsulates the claustral area where cell label is substantially attenuated. Thick dashed lines demarcate the border between the orbitofrontal and insular cortices, while the thin dashed lines designate the approximate borders of the claustrum at rostral levels. Ins, insular cortex; LO, lateral orbital cortex; PIR, piriform cortex. Scale bars = 200 μm.

At striatal levels, a band of retrogradely labelled cell bodies was present in the insular cortex surrounding the claustrum, both superficial, i.e. juxtaposed to the external capsule) and deep to the claustrum. Within the claustrum core, very few retrogradely labelled cell bodies were present, particularly at more septal/striatal levels. Anterior to the striatum, the distribution of cortical label outlined a region of attenuated label that closely matched that which was defined by the differential expression of cortical tracers, Gng2, parvalbumin and Crym (**Fig. 9**). In two of these cases, stained sections for parvalbumin confirmed that the region of attenuated label was indeed claustrum. In these same two cases overlays with cresyl stained section confirmed that the border between the lateral orbital and the insular cortices co-localise with the parvalbumin based definition of claustrum.

## Discussion

A consensus on the anatomical boundary of the claustrum-endopiriform complex is important for establishing its functional role and, on a more immediate and practical level, for both the interpretation of, e.g. anatomical studies, as well as in the verification of electrode placements in electrophysiological studies.

Our primary finding is that the expression profiles of three claustral marker genes, Gng2 (**Fig. 6**), parvalbumin (**Fig. 1**) and Crym (**Fig. 5**), as well as cortical (**Figs. 7–8**) and thalamic (**Fig. 9**) tracing data, demonstrate that the anatomical boundary of the rat claustrum extends approximately 500 μm rostral to the anterior horn of the neostriatum, remaining confined throughout its rostro-caudal span to layer 6 of the insular cortex (**Fig. 10**). Our findings relating to the *caudal* extent of the claustrum in the rat are in close accordance with atlas-based delineations (e.g. Paxinos and Watson, 2005), where vCLA and dCLA terminate at the level of the transition of insular to rhinal cortices (**Fig. 2**). Caudal to this coronal level, the claustral differential expression profiles of Crym, Gng2, and parvalbumin within the deep insular cortex were no longer apparent.

**Figure 10.**
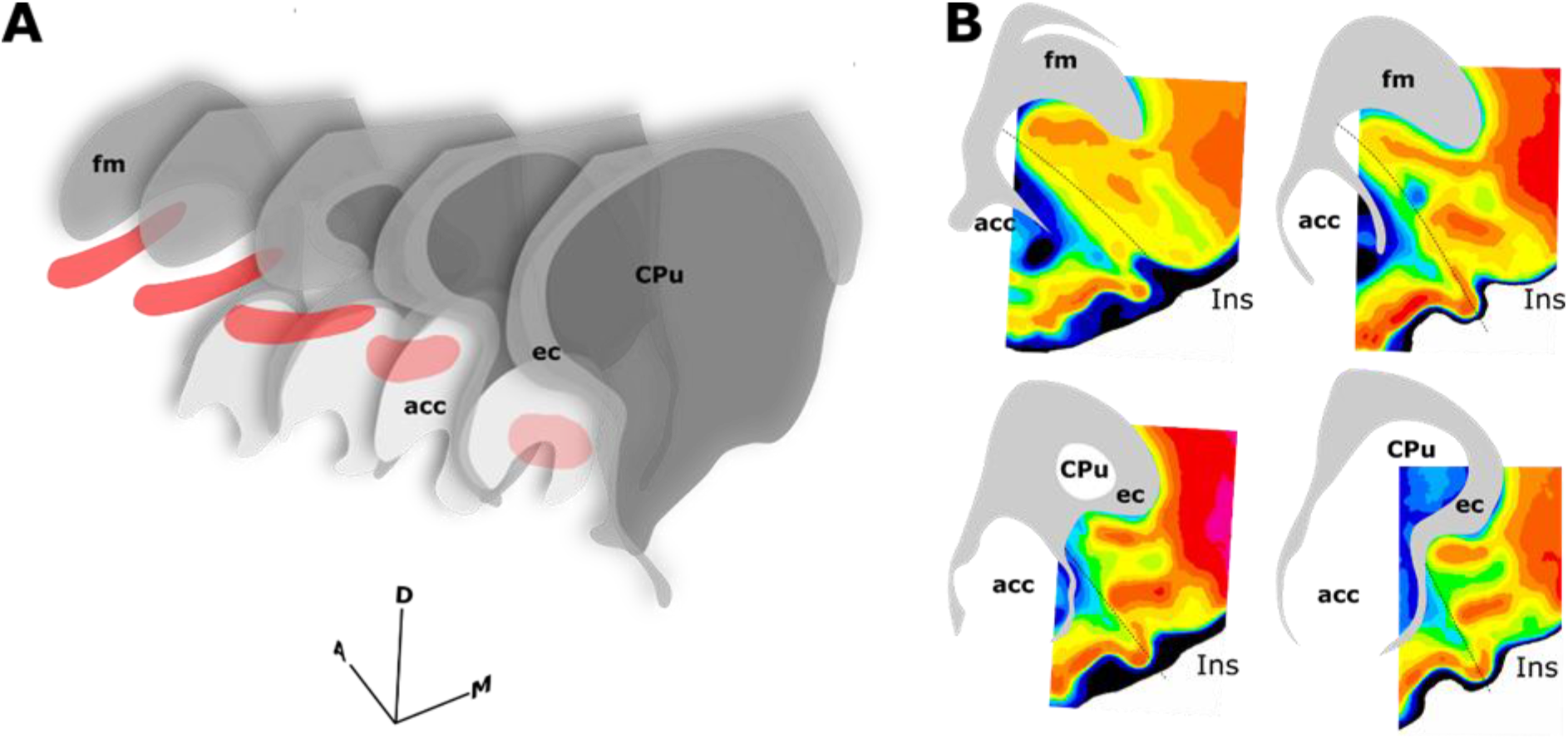
Schematics summarising the rostral extent of the anatomical boundary of the claustrum in the rat. Expression of parvalbumin, crystallin mu and Gng2, along with anterograde and retrograde tracing of claustro-cortical connectivity, and retrograde tracing of corticothalamic connectivity provided highly complimentary definitions of the boundary outlined in **A-B. A**, 3-dimensional representation of parvalbumin expression (red) in the ventral claustrum from caudal striatal (CPu) anterior-posterior levels (front) to rostral levels anterior to the striatum (back). Note the change in orientation of the long axis of the claustrum as it follows the arch of the forceps minor of the corpus callosum (fm) and the external capsule (ec). **B**, Pixel density-based heatmaps (warm colours represent high expression) of parvalbumin expression in sections anterior to the striatum (upper) and at the level of the striatum (lower), reinforcing the extent and continuity rostral to the striatum. Abbreviations: acc, nucleus accumbens; Ins, insular cortex.

In findings that are consistent with reports both in the mouse (Wang et al., 2017) and rat (Mathur et al., 2009), expression of Crym was dense in the insular cortex. At striatal levels, dense distributions of Crym-immunoreactive cell bodies and neuropil were distributed around the circumference of the claustrum, very clearly delineating the cortical shell surrounding the claustrum as described previously (Mathur et al., 2009). Within the core of the vCLA, Crym-immunoreactivity (both cell bodies and neuropil), was all but absent except for a handful of Crym-immunoreactive (putative) ectopic cortical cell bodies. Conversely, Gng2 and parvalbumin expression was enriched in the core of the vCLA and delineated a boundary that closely matched that which was negatively outlined by the Crym expression profile. Mathur and colleagues (2009) established, through dual-immunofluorescence, that parvalbumin expression in the claustrum revealed an anatomical boundary of the claustrum that closely matched that shown by Crym (**Fig. 10**). Using manual registration of serial sections, we have shown that Crym and Gng2 outline a similarly consistent nuclear boundary, providing further validation of these markers (Supplementary **Fig. S2**). Retrograde cortical and thalamic tracing experiments, as well as anterograde cortical tracing cases provided data that was highly complementary to our IHC findings. The rostral claustrum has been suggested to be positioned deep to the ventral and lateral orbital cortices (Paxinos and Watson, 2005). We observed that fibres from the anterior cingulate cortex terminated densely in this area but, importantly, that this fibre plexus did not co-localize with the parvalbumin label, thereby further consolidating the idea that this area is cortical and not claustral.

Numerous studies have reported dense connectivity between the claustrum and the thalamus in the rat (Herkenham, 1978a; McKenna and Vertes, 2004; Vertes et al., 2006, 2012; Yoshida et al., 2005; Zhang et al., 2001). Given the cortical shell surrounding the claustrum core and the presence of ectopic cortical neurons throughout its rostral-caudal extent (Mathur et al., 2009), there is uncertainty as to whether these reported connections are accurate. Our data seems to support the case presented initially by Mathur and colleagues (2009) that the cortex immediately surrounding the claustrum shares subcortical connectivity but the claustrum itself does not. Our retrograde thalamic injections included the three thalamic nuclei that have been suggested to receive claustral afferents (Erickson et al., 2004; Herkenham, 1978b; McKenna and Vertes, 2004; Vertes et al., 2012; Zhang et al., 2001) We did observe scattered thalamic projecting cells in the claustrum area, especially at rostral level. However, as the same pattern was seen in Crym stained sections, i.e. ectopic insular cortical neurons were occasionally scattered within the core of the claustrum, with the same rostro-caudal distribution, it is most likely that our retrograde labelled cells were in fact cortical cells.

In certain respects, our findings contradict those of Mathur and colleagues (2009), whose conclusions prompted a reassessment of the atlas-based anatomical boundary of the claustrum to one which: 1. Did not extend rostral to the anterior horn of the neostriatum, and 2. Was not juxtaposed to the external capsule but was instead surrounded by a cortical shell.

In their study, Mathur and colleagues also examined the expression of parvalbumin, Gng2, and Crym in the rat, using the same primary antibodies and similar dilutions. However, as most data were shown as immunofluorescence label, and not immunohistochemistry, it is possible that our immunohistochemical approach was more sensitive to identifying the slightly weaker frontal signal (in the Gng2 stain). Additionally, in their analysis of Gng2 and parvalbumin expression, photomicrographs depict an absence of label in the region ventral to the forceps minor of the corpus callosum, but one that is at an extreme rostral level in which this region is orbital, not insular (Mathur et al., 2009). The level depicted represents the rostral-most extent of the insular cortex at which level it is situated more laterally, i.e. outside of the presented field of view. In the same study, neuronal tract tracing was used in combination with parvalbumin immunofluorescent localisation and, in this instance, images were centered over parvalbumin immunofluorescence in the orbital cortices, in which no retrograde label was observed. It would, therefore, seem to be the case that Mathur and colleagues (2009) were correct in their disagreement with the atlas of Paxinos and Watson (2005), in that the claustrum is not situated within the orbital cortex at rostral levels, but mistaken in their conclusion that the claustrum was, therefore, only present at striatal levels. The consequence of these contradictory findings has been the development of a trend in many recent studies to include a methodological note stating that *analyses of claustral labelling did not extend beyond the most rostral coronal section that contained striatum due to the reported absence of Gng2 expression in these regions* contributing to an incomplete understanding of the claustrum.

As mentioned, the differential expression of Gng2 and Crym in the frontal extension of the claustrum becomes less accentuated. It would seem to be the case that towards the rostral apex of the claustrum, the density of ectopic cortical neurons within the core of the claustrum increases, constituting something of a claustro-cortical transition (**Fig. 4** and supplementary **Fig. 1**), but it is also worthy of note that at these rostral levels, ascending axon bundles from neurons within the insular/orbital cortices enter the forceps minor of the corpus callosum in a path that bisects the claustrum (Coizet et al., 2017). These bundles would appear to reduce both the uniformity of Crym attenuation and the clarity of the gene marker-defined boundary.

Parvalbumin expression in the rodent CLA-DEn complex is confined to the vCLA (Smith et al., 2018), avoiding the DEn and the dCLA. As a result, the distribution of parvalbumin provides an important reference in determining the extent of gng2 and Crym expression within the CLA-EN complex and, of relevance here, the relative components of the rostral extent of the complex. At the anterior horn of the neostriatum, insular and piriform cortices are juxtaposed with vCLA embedded within layer 6 of the insular cortex and DEn within the deepest layer 3 of the piriform cortex. At this level, vCLA and DEn are continuous. Further rostrally, the emergence of the orbital cortex separates insular and piriform cortices and, therefore, vCLA from DEn. Meanwhile, vCLA and dCLA remain continuous throughout the caudo-rostral extent of the complex. Rostral to the striatum, the vCLA/dCLA complex becomes situated progressively more ventrolaterally, relative to the forceps minor of the corpus callosum.

## Conclusions

Using neuroanatomical tracing and the expression profiles of two genes that are widely accepted to be differentially expressed in the striatal claustrum, we report here that, contrary to previous reports, the rostral extent of the claustrum in the rat extends anterior to the rostral apex of the striatum. Our combined tracing and gene-marker based data represent a unified view of the position of the rostral claustrum. The functions of claustrum are matter of continuing investigation, with cells that appear to code for aspects of extended space present in the rat claustrum (Jankowski and O’Mara, 2015), somewhat akin to the place cells and other spatial cells found in the hippocampal formation and other related areas (Grieves and Jeffery, 2015). The seeming absence of either thalamic or hippocampal inputs suggest that the spatial coding in claustrum observed by Jankowski and O’Mara (2015) is likely to be cortical in origin, perhaps originating from a combination of spatial inputs from, e.g., grid cells of the entorhinal cortex (Hafting et al., 2005), and other inputs from regions such as parieto-insular vestibular cortex (e.g. Rancz et al., 2015). The relative cortically-encapsulated inputs and outputs of claustrum we describe here would support this proposition.

### Author Contributions

CD, MM, SOM and MJ conceived and designed the experiments; CD, MM, BF, EB, ML performed the experiments; CD, MM, ML, SOM analysed the data; CD, MM wrote the manuscript with contributions from SOM, BF, MJ and JA; EB and JA contributed data or analysis tools.

### Funding

This work was supported by Science Foundation Ireland grant SFI 13/IA/2014 and a Joint Senior Investigator Award made by the Wellcome Trust to S.M.O.M. and J.P.A. #103722/Z14/Z

## Acknowledgements

The authors would like to thank Dr Lisa Kinnavane for her assistance with a retrograde tracing experiment.

## Supplementary Figures

**Figure S1.**
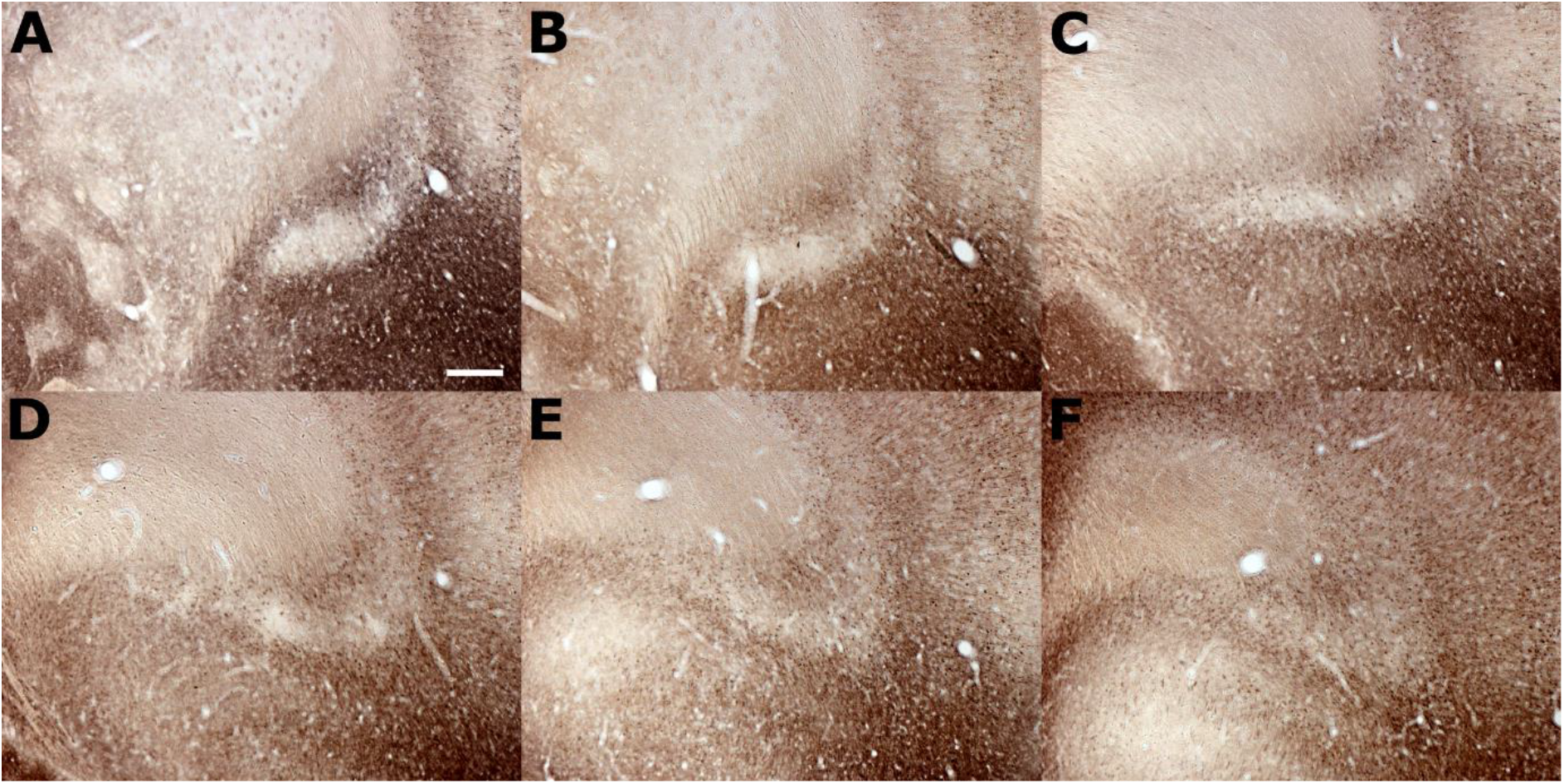
Photomicrographs of a sequential 1-in-4 series reacted against crystallin mu (Crym) ranging from a mid-striatal anterior-posterior level (A), to the rostral peak of the striatum (C) and up to a rostral aspect of the claustrum approximately 600 μm anterior to the striatum. Scale bar = 300 μm

**Figure S2.**
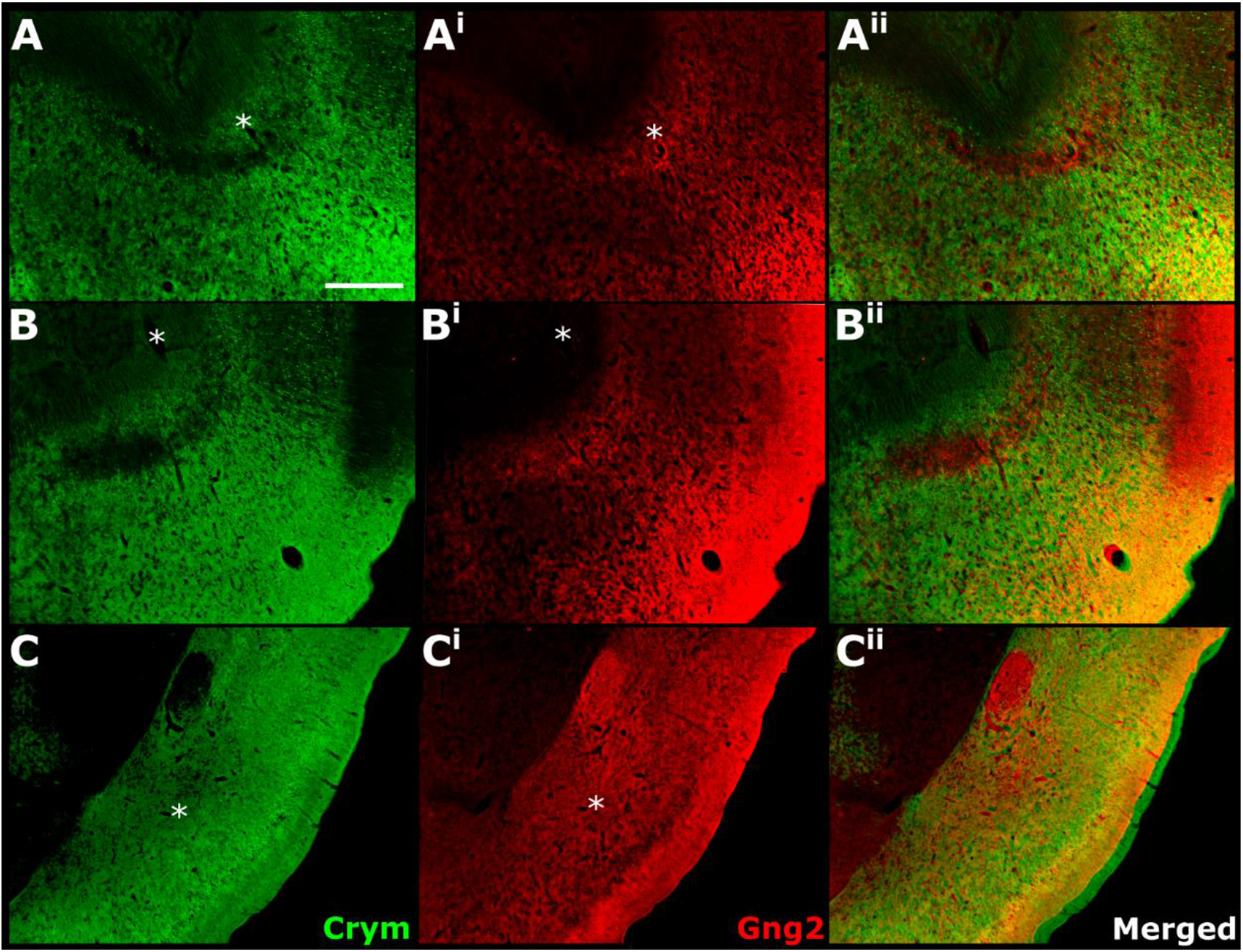
Images from sequential immunohistochemically (DAB) stained crystallin mu (Crym; **A-C**) and Gng2 (**A^i^-C^i^**) sections that have been converted to 8-bit and pseudo-coloured in green and red, respectively. Merged Images were aligned and manually registered using landmarks (asterisks) to assess the overlap between attenuated Crym staining in the claustrum and enriched Gng2 expression. **A-A^ii^** shows overlap between Crym and Gng2 rostral to the striatum; **B-B^ii^** shows overlap at the rostral apex of the striatum and **C-C^ii^** show overlap at a mid-striatal anterior posterior level. In all cases, Gng2 enrichment and Crym attenuation delineated a consistent claustrum border. Scale bar (applies to all) = 500 μm.

**Figure S3.**
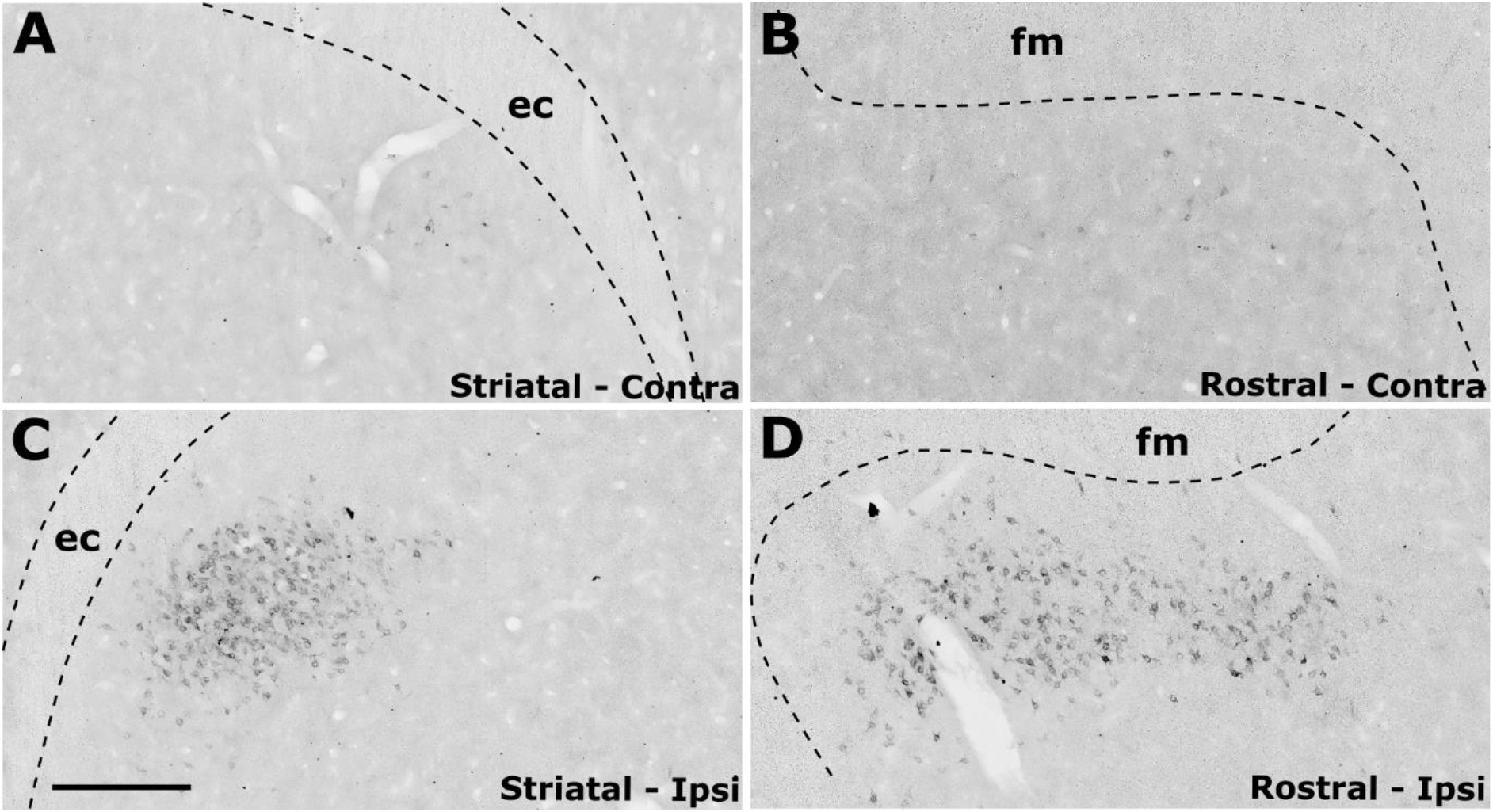
Unilateral (right hemisphere) retrograde Tracer injections (Fluoro-gold; FG) targeting the retrosplenial cortex resulted in labelled cell soma in the claustrum both at striatal anterior-posterior (AP) levels (**C-D**) as well as rostral to the striatum (**A-B**) in a distribution that closely matched parvalbumin expression in the claustrum (See dual fluorescent label (FG and parvalbumin) from the same case in **Fig. 8**). Scale bar (applies to all) = 300 μm.

**Figure S4.**
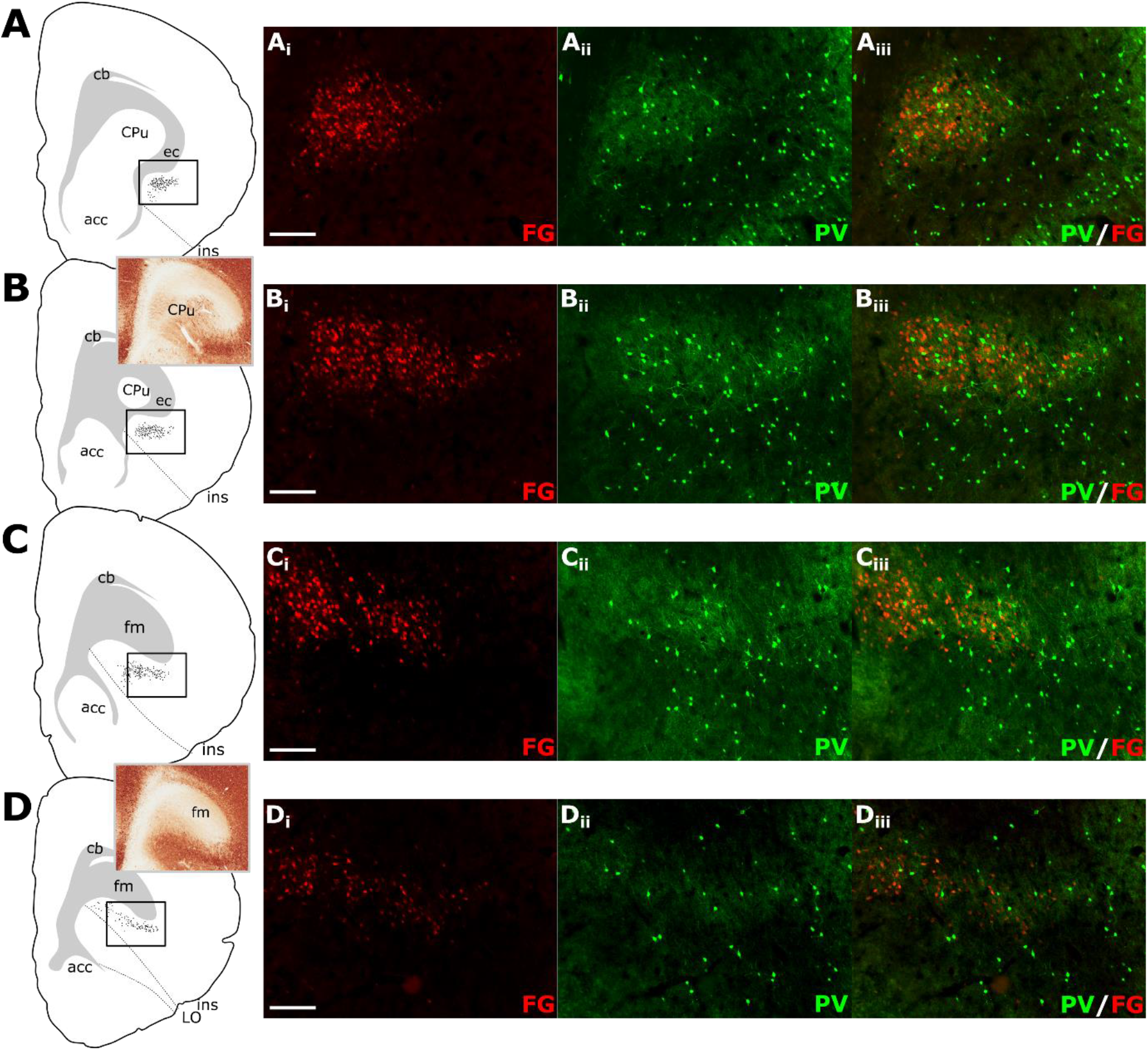
Tracer injections of Fluorogold (FG) within retrosplenial cortex resulted in dense retrograde label throughout the extent of the ipsilateral claustrum. **A-D**: Schematic tracings of caudal (striatal (CPu); **A-B**) to rostral (anterior to striatum; **C-D**) brain sections showing retrograde label in the claustrum*. Rectangles in **A-D** show regions shown in corresponding fluorescence micrographs (i-iii). Dual-fluorescence experiments showed that parvalbumin neuropil expression (PV; green) closely overlaid that of the FG retrograde label (FG; red). Insets in **B** and **D** show anterior-posterior level relative to CPu in parvalbumin-reacted tissue. Scale bars = 200 μm.

**Figure S5.**
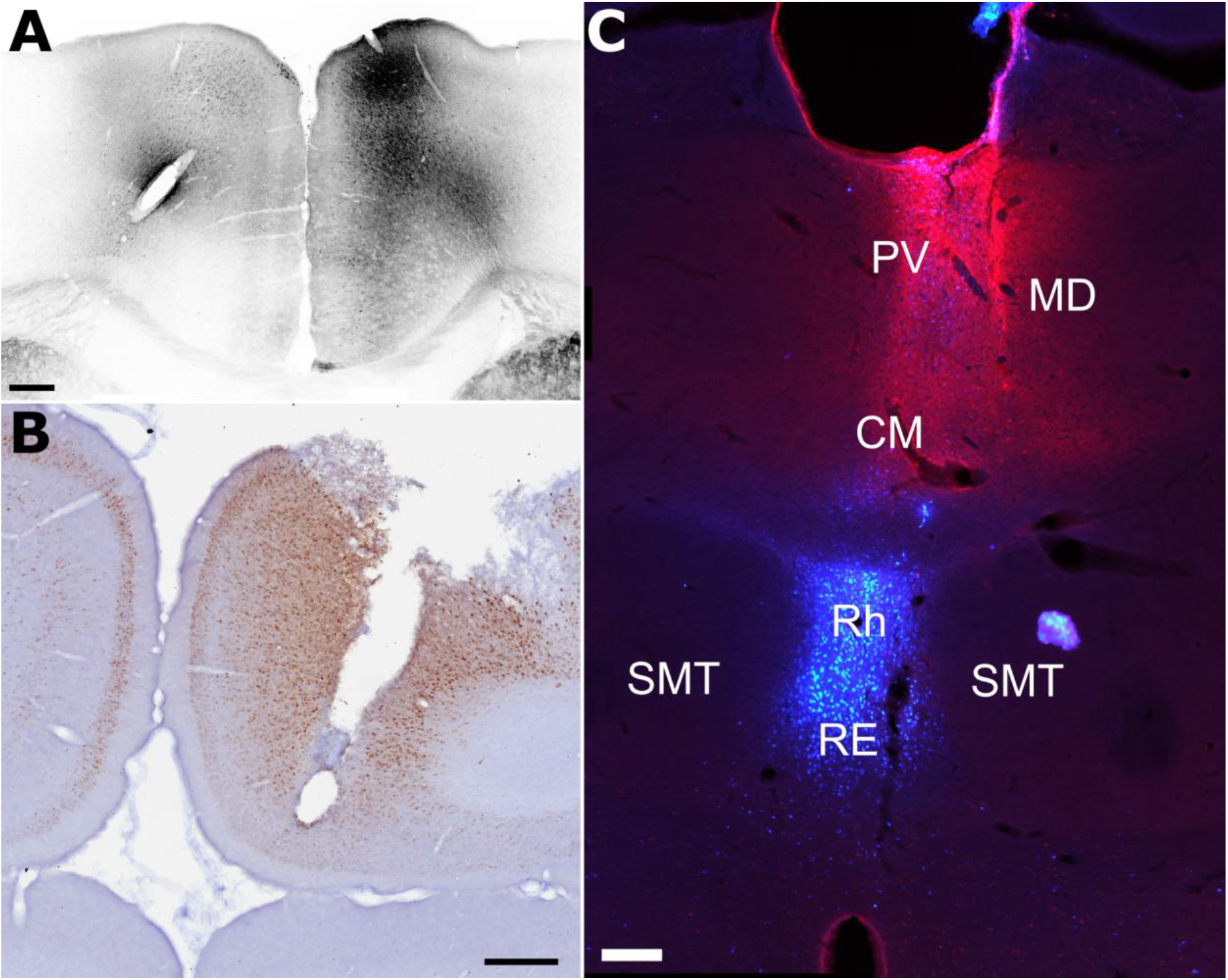
Cortical (**A-B**) and thalamic (**C**) pressure injections of neuronal tracer were used to assess claustrum connectivity profiles. **A**, an example of a pAAV-CaMKIIa-hM4D(Gi)-Mcherry pressure injection into the anterior cingulate cortex (case 219#3); **B**, an example of a Flouro-gold pressure injection into the anterior cingulate cortex (FGRSC1) **C**, An example of an injection site of cholera-toxin b (red) and Fast Blue (blue) injections sites in the centromedial (CM)/paraventricular (PV)/mediodorsal (MD) and nucleus reuniens (RE)/rhomboid (Rh), respectively. Abbreviations: SMT, submedius thalamic nucleus. Scale bar in **A** = 400 m; **B-C** = 200 μm.

